# High-resolution interrogation of functional elements in the noncoding genome

**DOI:** 10.1101/049130

**Authors:** Neville E. Sanjana, Jason Wright, Kaijie Zheng, Ophir Shalem, Pierre Fontanillas, Julia Joung, Christine Cheng, Aviv Regev, Feng Zhang

## Abstract

The noncoding genome plays a major role in gene regulation and disease yet we lack tools for rapid identification and manipulation of noncoding elements. Here, we develop a large-scale CRISPR screen employing ~18,000 sgRNAs targeting >700 kb of noncoding sequence in an unbiased manner surrounding three genes (*NF1, NF2*, and *CUL3*) involved in resistance to the BRAF inhibitor vemurafenib in the *BRAF*-mutant melanoma cell line A375. We identify specific noncoding locations near genes that modulate drug resistance when mutated. These sites have predictive hallmarks of noncoding function, such as physical interaction with gene promoters, evolutionary conservation and tissue-specific chromatin accessibility. At a subset of identified elements at the *CUL3* locus, we show that engineered mutations lead to a loss of gene expression associated with changes in transcription factor occupancy and in long-range and local epigenetic environments, implicating these sites in gene regulation and chemotherapeutic resistance. This demonstration of an unbiased mutagenesis screen across large noncoding regions expands the potential of pooled CRISPR screens for fundamental genomic discovery and for elucidating biologically relevant mechanisms of gene regulation.

---

More than 98% of the human genome is noncoding, however, unlike the coding genome there exists no overarching theoretical framework (e.g. protein triplet code) capable of translating noncoding genomic sequence into functional elements (Maher 2012; Visel et al. 2009). Evidence from genome-wide association studies (GWAS) suggests many noncoding regions are critical for human health and disease: more than 2600 single-nucleotide polymorphisms (SNPs) have been associated with human disease/traits, the vast majority (>97%) of which occupy noncoding regions (Hindorff et al. 2009; Maurano et al. 2012; Schaub et al. 2012). The significance of these associations, however, has been difficult to assess, in part because we lack the tools to determine which variants alter functional elements. In recent years, there have been major advances in identifying molecular hallmarks that correlate with putative functional elements in the noncoding genome, such as epigenetic state, chromatin accessibility, transcription factor binding, and evolutionary conservation. Consortium efforts such as the Encyclopedia of DNA Elements (ENCODE) and the Roadmap Epigenomics project have produced a vast amount of genome-scale data that is widely used to predict regulatory function (Maher 2012; Roadmap Epigenomics Consortium et al. 2015). However, these predictions largely bypass regions for which there are no hallmarks, and it is difficult to ascertain if these hallmarks play a correlative or truly causal role in function or phenotype (Kwasnieski et al. 2014; Mundade et al. 2014). Experimental efforts to determine causality have employed episomal reporters that utilize preselected DNA fragments with expression serving as a proxy for function (Melnikov et al. 2012). These methods assess the DNA fragments in plasmids and are therefore decoupled from the local chromatin context and broader regulatory interactions, both of which are important characteristics of gene regulatory mechanisms. Thus there is a need for systematic approaches to sift through noncoding variants and determine if and how they affect phenotypes within a native biological context.

To address this need, we designed a high-throughput method using pooled CRISPR (Clustered regularly-interspaced short palindromic repeat)-Cas9 libraries to screen noncoding genomic loci to identify functional regions related to phenotype and gene regulation. Previous applications of CRISPR screens within the noncoding genome have focused on select elements, such as miRNAs, enhancers based on predictions derived from chromatin immunoprecipitation (ChIP) of functional hallmarks, or transcription factor binding, but they have not gone beyond these sequences (Chen et al. 2015; Canver et al. 2015; Diao et al. 2016; Korkmaz et al. 2016). Here, we comprehensively assay a total of 715 kb of sequence surrounding three different genes by performing unbiased mutagenesis to uncover functional elements relevant to cancer drug resistance. This approach requires no pre-existing knowledge of the region being screened and enables discovery of both gene-proximal and gene-distal functional elements.

Vemurafenib is a potent inhibitor of mutant BRAF, which is found in 50-70% of melanomas (Hodis et al. 2012; Cancer Genome Atlas Network 2015). Resistance to vemurafenib arises within months in almost all patients (Sosman et al. 2012) and surviving tumor cells display increased malignancy that rapidly leads to lethality (Zubrilov et al. 2015). Previously, we used a genome-scale CRISPR library to identify genes in which loss-of-function mutations result in resistance to vemurafenib in a melanoma cell line with a V600E BRAF mutation (Shalem et al. 2014). To explore if mutations in the noncoding regions around three of the previously validated resistance genes (*NF1*, *NF2*, and *CUL3*) could similarly impact drug resistance, we designed three single-guide RNA (sgRNA) libraries tiling across 100 kb regions 5´ and 3´ of each gene (Fig. 1A). For each library, we synthesized the sgRNAs as a pool (6,682 for *NF1*, 6,934 for *NF2*, and 4,699 for *CUL3;* 18,315 sgRNAs total) and cloned them into a lentiviral vector (fig. S1A, B). Using the A375 BRAF V600E human melanoma cell line expressing Cas9, we transduced cells with these pooled sgRNA libraries at a low multiplicity of infection (~0.2 virions/cell) and selected for cells that received a sgRNA (Sanjana et al. 2014). After 7 days of selection (and Cas9-mediated genome modification), A375 cells were cultured in 2uM vemurafenib or control (DMSO) for 14 days. Using deep sequencing, we counted the representation of sgRNAs in the library in each condition and compared it with an early time point taken immediately before the drug/control treatment (Fig. 1B-D, *left*). Compared to this early time point, control cells had minimal changes in library representation, whereas cells treated with vemurafenib showed greater variability in sgRNA representation. We fit a linear model to the control distribution to detect enriched sgRNAs in vemurafenib-treated cells (enriched >4 standard deviations from the control distribution), which we displayed as a function of genomic coordinates in a genome browser-style view (Fig. 1B-D, *right*). An enriched sgRNA suggests that the sgRNA target site may contain a functional noncoding sequence that increases vemurafenib resistance and improves the survival of A375 cells.

**Figure 1.**
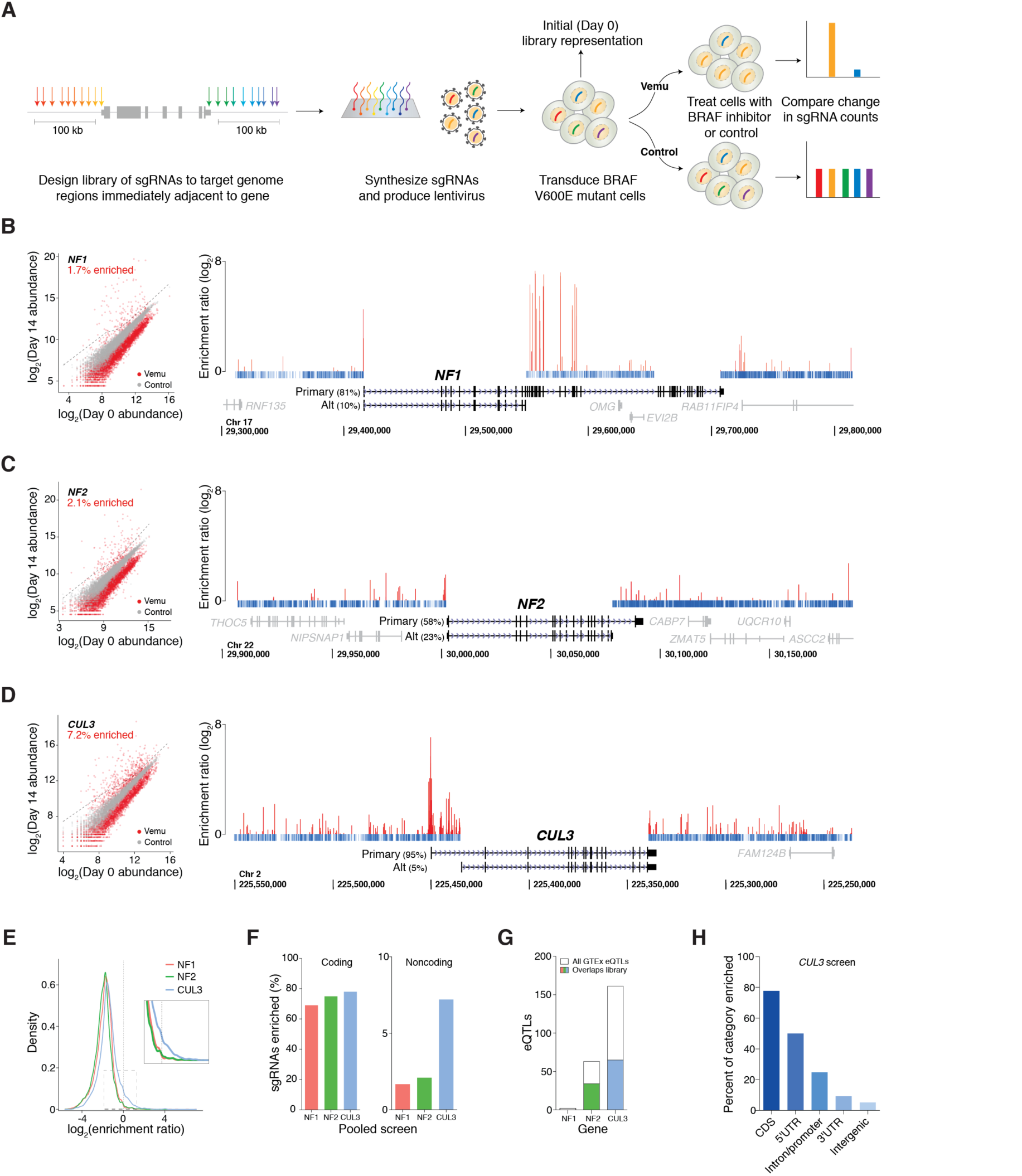
CRISPR mutagenesis of ~200 kb noncoding regions flanking three genes involved in BRAF inhibitor resistance. A) Design of sgRNA libraries targeting 100 kb 5` and 100 kb 3` of a gene locus. After library design, sgRNAs are synthesized on an array and cloned into a lentiviral vector. BRAF mutant cells are transduced with the pooled lentivirus and treated with control (DMSO) or the BRAF inhibitor vemurafenib (vemu) for 14 days. Using a deep sequencing readout, sgRNAs that are enriched after treatment with vemurafenib are identified by comparison with an early time point (Day 0) and cells treated with control. B) - D) (*left*) Scatterplot of normalized read counts for each sgRNA at Day 0 (*x* axis) and at Day 14 (*y* axis) for 3 mutagenesis screens (B: *NF1*, C: *NF2*, D: *CUL3*). Gray dots indicate read counts from control cells and red dots indicate read counts from vemurafenib-treated cells. Dotted line denotes 4 standard deviations from the mean of the control cell distribution. The percentage of enriched sgRNAs in vemurafenib (>4 s.d.) is shown. (*right*) Enrichment ratio for 3 separate mutagenesis screens targeting ~200 kb near gene loci (B: *NF1*, C: *NF2*, D: *CUL3*) in A375 *BRAF* mutant cells. sgRNAs are plotted by genome coordinates (hg19) of their target site. The enrichment ratio is the log_2_ ratio of the normalized read count for each sgRNA in vemurafenib to its normalized read count in control (minimum from 2 replicate screens). Enriched sgRNAs are plotted in *red* with their enrichment ratio. For depleted sgRNAs (*blue*), only position is shown. Relative expression from RNA-seq in A375 of the top two RefSeq isoforms for each gene is indicated next to the corresponding transcript. All gene-specific libraries were designed to target the proximal 100 kb from the start/end of each RefSeq isoform’s coding sequence. E) Distribution of log_2_ ratio of the normalized read count for each sgRNA in vemurafenib to its normalized read count in control (minimum over 2 replicate screens). F) Percent of sgRNAs that are enriched (>4 s.d. from control cells) with target sites in coding regions (*left*) or noncoding regions (*right*) for the *NF1*, *NF2*, and *CUL3* pooled screens. G) Total expression quantitative trait loci (eQTLs) found in the Genotype-Tissue Expression (GTEx) v6 analysis release (7,051 tissue samples from 449 donors) for *NF1*, *NF2*, and *CUL3*. Shaded regions indicate eQTLs that are contained within the region targeted by each sgRNA library. H) Percent of enriched sgRNAs by genomic category (coding sequence [CDS], 5` UTR, promoter/first intron, 3` UTR, and intergenic) in day 14 vemurafenib-treated cells.

Overall, most sgRNAs were depleted after treatment with vemurafenib, which is expected since vemurafenib targets the oncogene addiction that drives A375 growth (Fig. 1E). However, in all three libraries, we found a small group of sgRNAs that were enriched after vemurafenib treatment (log_2_ ratio of Vemu/Control > 0), with the *CUL3* library having the largest percentage of enriched sgRNAs. In our library design, we also included a small number of sgRNAs targeting the coding region of each gene and, as expected, most sgRNAs targeting coding regions (70 - 80%) were enriched for each gene. However, amongst the sgRNAs targeting noncoding regions, approximately 4-fold more sgRNAs were enriched in the *CUL3* library than in the *NF1* or *NF2* libraries (7.2% of noncoding sgRNAs in the *CUL3* library, 1.7% in the *NF1* library, and 2.1% in the *NF2* library), suggesting the presence of more gene regulatory elements in the noncoding regions flanking the gene (Fig. 1F). To determine if this increase in putative gene regulatory elements in the 200 kb region surrounding *CUL3* is also reflected in human gene expression and genotyping data, we queried expression array and RNA sequencing data from the Genotype-Tissue Expression (GTEx) database v6 (7,051 tissue samples from 449 donors).

Indeed, we found that *CUL3* had the largest number of *cis-*expression quantitative trait loci (eQTL) (*n* = 161 eQTLs, mean effect size = ∐0.21), and the region targeted by the sgRNA library overlaps with a large number of these eQTLs (Fig. 1G) (GTEx Consortium 2015). Given the relatively greater number of putative regulatory elements from our CRISPR screen and from the GTEx data, we chose to focus our downstream analysis and validation efforts on *CUL3*. Among noncoding regions targeted in the *CUL3* library, we found that a higher percentage of sgRNAs targeting gene-proximal elements were enriched compared to other noncoding regions (Fig. 1H) and, in general, we observed greater enrichment for sgRNAs targeting noncoding elements on the 5´ side of the gene (e.g. promoter, 5´ untranslated region [UTR]) than for those on the 3´ side (fig. S1C).

To understand the distribution of enriched sgRNAs from the *CUL3* locus, we designed multiple analyses to identify the properties of the enriched sgRNA target sites. One method by which distal elements can regulate gene expression is through interactions with the promoter region. This can occur due to chromatin looping and close proximity between regions in three dimensions despite large (linear) distances (Lieberman-Aiden et al. 2009). To test if regions targeted by enriched sgRNAs from the screen physically interact with the *CUL3* promoter, we created three independent chromosome conformation capture (3C) libraries to test for interactions over the screened region with the *CUL3* promoter (Fig. 2A) (Dekker et al. 2002; Miele et al. 2006). We designed droplet digital PCR (ddPCR) probe combinations to quantify the interaction frequency for each potential interacting site across the ~200 kb region. In total, the interaction frequencies of 156 possible interactions with the *CUL3* promoter region were measured (table S1). We found that regions on the 5´ side of *CUL3* tend to interact more strongly with the promoter (in agreement with greater sgRNA enrichment on the 5´ side) and that regions with higher 3C interaction contain, on average, more vemurafenib-enriched sgRNAs (Fig. 2B).

**Figure 2:**
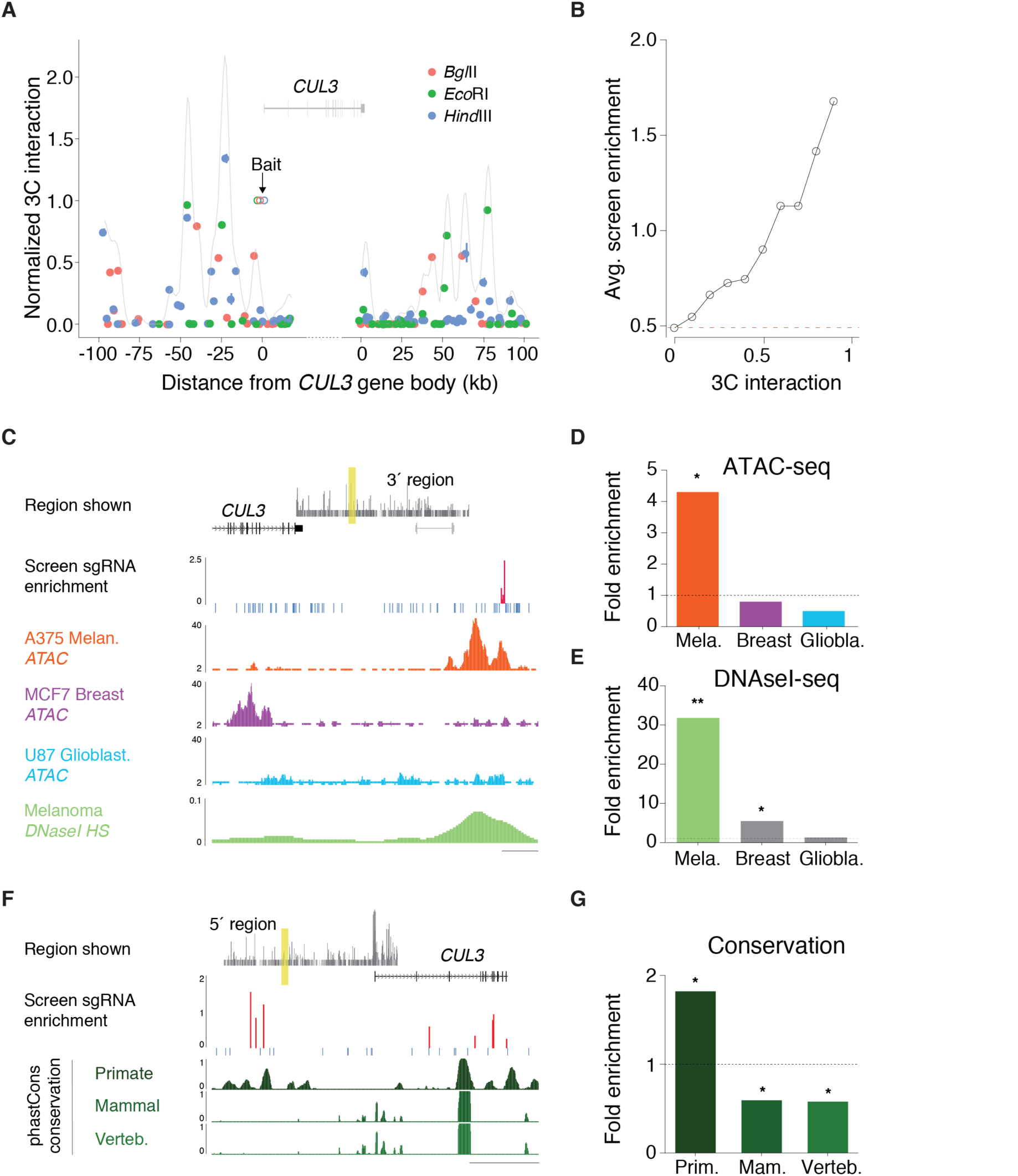
Functional noncoding elements at the *CUL3* locus correlate with physical chromatin interactions, chromatin accessibility and recent evolutionary conservation. A) Plot of interaction frequencies with the *CUL3* promoter based on chromatin conformation capture (3C) in A375 cells. Data points represent three independent 3C libraries generated with three separate restriction enzymes (*BglII, EcoRI*, and *HindIII*). The grey curve shows a smoothed estimate of interaction frequency by convolution of the 3C data points with a Gaussian kernel. For the Gaussian kernel, the standard deviation is half the average distance between restriction sites in each library (4.3 kb). B) The average enrichment of sgRNAs (log_2_ ratio of vemurafenib/DMSO reads) near all 3C sites with an interaction frequency with the *CUL3* promoter equal to or greater than the indicated value. Nearby sgRNAs were grouped into overlapping windows of the same size as the average distance between restriction sites in each library (4.3 kb) and the closest window was selected for each 3C site. C) An example of enriched sgRNAs (*red*) that overlap with a melanoma-specific region of open chromatin. Assay for Transposable and Accesible Chromatin Sequencing (ATAC-seq) in A375 melanoma (*orange*), MCF-7 breast cancer (*purple*) and U-87 glioblastoma (*blue*) and Melanoma DNAse I hypersensitivity sequencing (DNAse I HS-seq) (*green*, ENCODE/OpenChromatin/Duke Colo-829). Approximate location of region (3´ of *CUL3*) is shown at top(*yellow* highlighted region). Scale bar: 500 bp. D) Fold enrichment of enriched sgRNAs near ATAC-seq open chromatin peaks in melanoma, breast cancer and glioblastoma cell lines. Fold-enrichment is computed by first finding the average sgRNA enrichment near ATAC peaks over the entire region targeted by the sgRNA library. This quantity is then divided by the mean of a distribution of the same quantity calculated from 10,000 random reshufflings of open chromatin peaks. E) Fold enrichment of enriched sgRNAs near DNAse I HS-seq (*below*) open chromatin peaks in melanoma, breast cancer and glioblastoma cell lines. Fold-enrichment is computed by first finding the average sgRNA enrichment near DNAse peaks over the entire region targeted by the sgRNA library. This quantity is then divided by the mean of a distribution of the same quantity calculated from 10,000 random reshufflings of open chromatin peaks. DNAse I HS data is from ENCODE/OpenChromatin/Duke. F) An example of enriched sgRNAs (red) that coincide with regions that show primate-specific conservation. Primate, placental mammal and vertebrate conservation represented as phastCons probabilities (two-state phylogenetic hidden Markov model). Approximate location of region (5´ of *CUL3*) is shown at top (*yellow* highlighted region). Scale bar: 200 bp. G) Fold enrichment of enriched sgRNAs near phastCons (conserved sequence) peaks in primates, placental mammals and vertebrates. Fold-enrichment is computed by first finding the average sgRNA enrichment near phastCons peaks over the entire region targeted by the sgRNA library. This quantity is then divided by the mean of a distribution of the same quantity calculated from 10,000 random reshufflings of phastCons peaks.

In addition to physical interactions, chromatin accessibility is often used to identify regulatory elements (Crawford et al. 2004; Buenrostro et al. 2013). To quantify chromatin accessibility, we performed Assay of Transposase-Accessible Chromatin with high-throughput sequencing (ATAC-seq) using A375 melanoma cells and two human cancer cell lines that originate from different tissues: MCF7 breast cancer (lung metastasis to breast) and U87 glioblastoma. We also examined available DNase I hypersensitivity with high-throughput sequencing (DNase-seq) data from ENCODE for similar cell lines. We identified regions with enriched sgRNAs that overlapped with A375-specific ATAC-seq peaks and melanoma-specific DNase-seq peaks (Fig. 2C) and, overall, we found higher sgRNA enrichment near A375-specific ATAC and melanoma-specific DNAse peaks than with chromatin accessibility from other cell types (Fig. 2D, E and fig. S2). This indicates that regions with enriched sgRNAs correlate with melanoma-specific open chromatin and may contain cell type-specific enhancers, consistent with previous results showing that enhancer histone marks are specific to particular cell or tissue types (Heintzman et al. 2007; Heintzman et al. 2009; Ernst et al. 2011; Sheffield et al. 2013).

A major hallmark of functional genome elements is evolutionary conservation of DNA sequence. As conservation varies widely across the noncoding genome, we tested whether more conserved regions harbor more enriched sgRNAs than less conserved regions. We examined phastCons conservation scores among primates (*n*=10 animals), placental mammals (*n*=33), and vertebrates (*n*=46) in the *CUL3* locus (Fig. 2F) (Felsenstein & Churchill 1996). Overall, enriched sgRNAs are ~1.8-fold more likely to be found near peaks of primate conservation and are ~1.7-fold less likely to be found near conservation peaks among mammals and vertebrates (Fig. 2G and fig. S2). In contrast, the genomic sites of sgRNAs targeting coding regions of *CUL3* do not demonstrate differential conservation (phastCons probability ~ 0.95 in primates, mammals and vertebrates). Although the magnitudes of the effects are smaller than those with chromatin accessibility, enriched noncoding sgRNAs preferentially target genomic regions that are more recently conserved (e.g. in primates) versus those conserved over longer evolutionary timescales.

Although these properties of enriched sgRNA target sites suggest functionality, we wanted to confirm that mutations in these specific noncoding regions lead to altered drug resistance and to test if these changes were mediated by *CUL3*. To assay specific sites for noncoding function, we individually cloned 25 sgRNAs that had a positive enrichment ratio into lentiviral vectors and produced virus (Fig. 3A and table S2). For this validation set, we selected sgRNAs that have at least one other similarly enriched sgRNA within 500 bp. We also attempted to choose these groups of sgRNAs for our validation set from several different genomic regions (e.g. 5´ and 3´ UTRs, promoter, intron, distal 5´ and 3´ regions) in order to understand the relative regulatory ability of noncoding elements across different locations. We transduced each lentivirus individually into A375 cells. After selection for 7 days, we amplified genomic DNA regions surrounding each sgRNA target and found an average of 85% of amplicons contained insertion-deletion (indel) mutations with near complete genome editing at most target sites (mean deletion size = 11 bp, mean insertion size = 4 bp, *n >* 5000 reads per site) (fig. S3) (table S3). After verifying genome modification at the targeted sites, we measured *CUL3* expression using a sensitive ddPCR hydrolysis probe assay. We found that 24 out of the 25 validation sgRNAs resulted in decreased *CUL3* expression relative to non-targeting sgRNAs (Fig. 3B, *left*). As expected, sgRNAs that target coding exons of *CUL3* resulted in an even greater loss of *CUL3* expression. We also treated cells transduced with sgRNAs from out validation set with 2uM vemurafenib and measured cell survival (vemurafenib resistance) individually: As expected, there is a negative correlation between *CUL3* gene expression and vemurafenib resistance (*r* = ∐0.54, *p* = 0.005, correlation does not include non-targeting sgRNAs or sgRNAs that target *CUL3* coding exons) (Fig. 3B, *right*). As a group, the validation sgRNAs targeting noncoding regions around *CUL3* produce moderate decreases in *CUL3* expression, which result in moderate increases in vemurafenib resistance.

**Figure 3:**
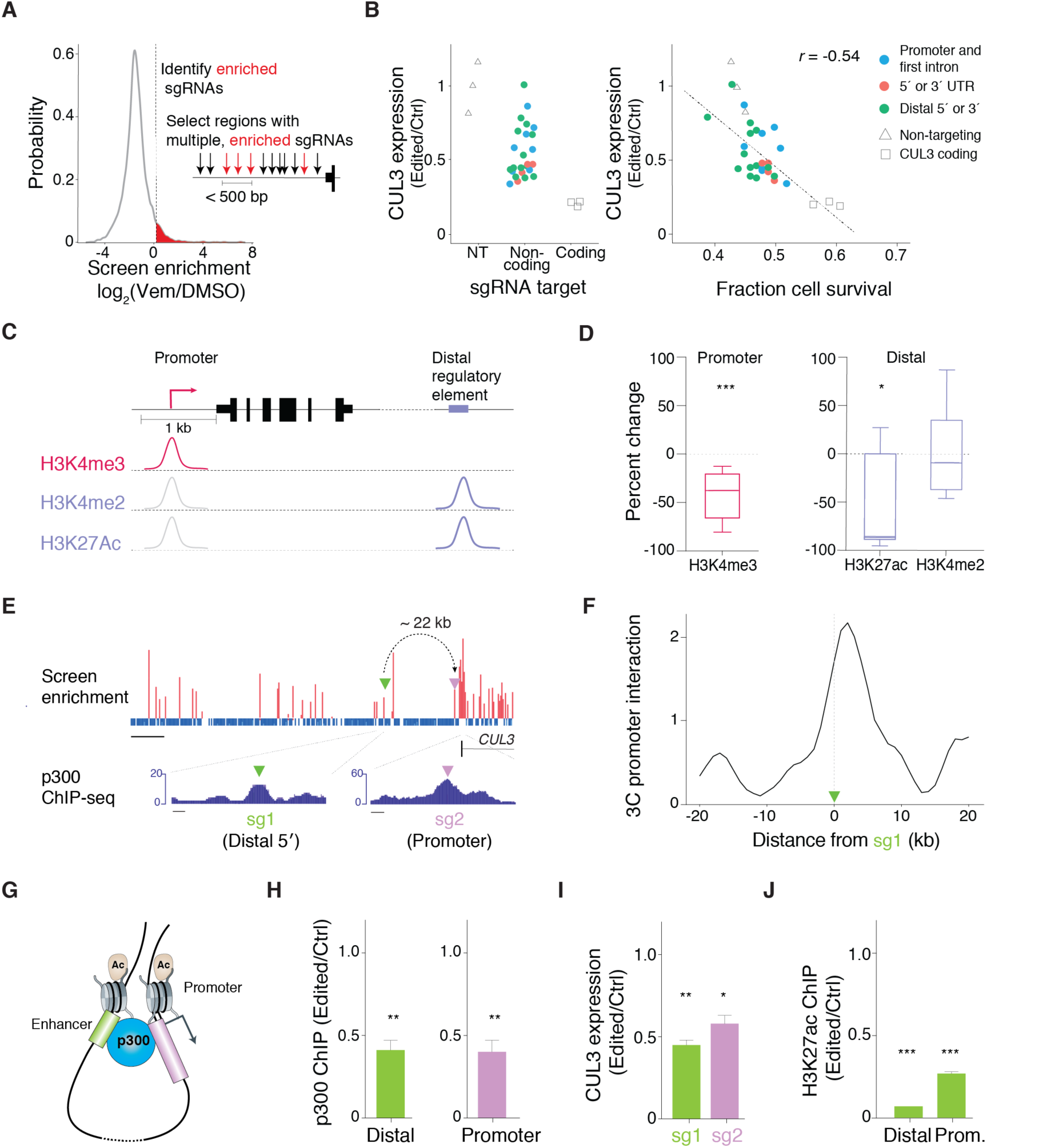
Noncoding mutations impact *CUL3* expression via long-range and local changes to the epigenetic landscape. A) Criteria for selection of a subset of library sgRNAs targeting noncoding regions for individual cloning and validation. The sgRNAs chosen for follow-up validation are enriched (log2 ratio of normalized vemurafenib/DMSO read counts > 0) and have at least one other similarly enriched sgRNA within 500 bp. From this group, a subset of 25 sgRNAs across the diversity of genomic categories (CDS, 5` UTR, promoter/first intron, 3` UTR, neighboring gene exon, and intergenic) was chosen for follow up studies. B) (*left*) *CUL3* RNA expression in A375 cells after transduction with lentivirus carrying non-targeting (*triangles*), selected noncoding region-targeting (*colored circles*) and exon-targeting (*squares*) sgRNAs. Changes in *CUL3* mRNA were quantified using droplet digital PCR (ddPCR) and all values are normalized to the median of cells transduced with non-targeting sgRNAs. (*right*) Relationship between *CUL3* expression and cell survival in A375 cells after 3 days of treatment with 2uM vemurafenib. Cells were transduced with lentivirus carrying non-targeting (*triangles*), selected noncoding region-targeting (*colored circles*) and exon-targeting (*squares*) sgRNAs. Linear fit and correlation is only to noncoding sgRNAs (*r* = ∐0.54, *p* = 0.005) and does not include exon-targeting or non-targeting sgRNAs. C) Schematic of histone modifications typically found at promoter proximal and distal regulatory elements. H3K4me3 is often found at the transcription start site of active or poised genes, whereas H3K27ac and H3K4me2 are found both at promoters and distal regulatory elements. D) Percent change in average H3K4me3 chromatin immunoprecipitation (ChIP) at 7 days post-transduction for all validation sgRNAs within 1 kb of the transcription start site of *CUL3*. Percent change in average H3K27ac and average H3K4me2 chromatin immunoprecipitation (ChIP) at 7 days post-transduction for all validation sgRNAs outside of the promoter proximal region of *CUL3*. E) Screen enrichment near a promoter proximal and a distal sgRNA site that coincide with p300 ChIP-seq peaks (ENCODE/SYDH/p300). Dashed arrow indicates a strong interaction frequency measured between the distal site and the *CUL3* promoter by 3C. Scale bars: 10 kb (screen enrichment), 250 bp (p300 ChIP-seq). F) Smoothed 3C signal measuring *CUL3* promoter interaction around distal sgRNA site in (E). G) Model of chromatin looping interaction to bring p300 enhancer element into proximity with the *CUL3* promoter. H) p300 ChIP around cut sites at 7 days post-transduction with distal element-targeting or promoter-targeting sgRNA (normalized to cells transduced with non-targeting sgRNA). I) H3K27ac ChIP at promoter-proximal and distal sites at 7 days post-transduction with distal element-targeting sgRNA (normalized to cells transduced with a non-targeting sgRNA). J) *CUL3* expression at 7 days post-transduction with distal element- and promoter-targeting sgRNA (normalized to cells transduced with non-targeting sgRNAs).

To understand the mechanism by which mutations in the noncoding region reduce *CUL3* expression, we surveyed changes in post-translational histone modifications at these sites. We divided our validation set of noncoding sgRNAs into two categories: sgRNAs that target within 1 kb of the *CUL3* coding region (“promoter”) and those outside this region (“distal regulatory”) (Heintzman et al. 2007; Heintzman et al. 2009). At most promoters, lysine 4 of histone H3 is tri-methylated (H3K4me3) and marks transcription start sites of genes that are active or poised (Santos-Rosa et al. 2002). At active enhancer elements, there is increased acetylation of lysine 27 of histone H3 (H3K27Ac) (Creyghton et al. 2010) and di-methylation of H3K4 (H3K4me2) without enrichment of H3K4me3 (Heintzman et al. 2007) (Fig. 3C). For sgRNAs within 1 kb of the transcription start site of the primary *CUL3* isoform, we performed chromatin immunoprecipitation followed by ddPCR (ChIP-ddPCR) and quantified the enrichment of H3K4me3 (table S4). We found a 56% decrease, on average, of H3K4me3 levels after editing (*p* = 7x10^∐4^, *n* = 9 edited sites) (Fig. 3D), consistent with the reduced gene expression. At distal regulatory sgRNAs target sites, we quantified changes in H3K27ac and H3K4me2 using ChIP-ddPCR, finding a 41% decrease, on average, in H3K27ac (*p* = 0.02, *n* = 7 edited sites) after editing and no significant change in H3K4me2 (*p* = 0.82, *n* = 7 edited sites) (Fig. 3D), although a subset of these sites did show a decrease in H3K4me2 levels after editing (fig. S4A).

Given the observed changes in *CUL3* expression and the surrounding epigenetic environment, we explored the impact of noncoding mutagenesis on histone-modifying protein occupancy and activity. Two sites targeted by validation sgRNAs occupy local peaks of enrichment for a histone acetyl-transferase and transcriptional co-activator, p300 (Fig. 3E). p300 expression and localization is prognostic in *BRAF* mutant melanoma (Bhandaru et al. 2014), and histone deacetylase inhibitors have been shown to work synergistically with vemurafenib to treat cancer (Lai et al. 2013). Although the two p300 sites are separated by ~22 kb, our 3C data indicates a strong interaction (Fig. 3F) that could bring the distal p300 site close to the proximal p300 site, which overlaps with the promoter region of *CUL3* (Fig. 3G). To explore if sgRNAs targeting these p300 sites alter occupancy and acetylation, we performed ChIP-ddPCR at both sites using antibodies for p300 and H3K27ac. After genome modification with the respective sgRNAs, we found a ~50% loss of p300 occupancy at each site (Fig. 3H) and a similar decrease in *CUL3* expression (Fig. 3I). In addition, after editing at the distal site, we detect a 93% loss of H3K27ac at that site (Fig. 3J) while levels of H3K27ac at a positive control region distant from the *CUL3* locus were unchanged (fig. S4B). Furthermore, we find a 75% decline in H3K27ac at the promoter site after editing at the distal site (Fig. 3J). These findings suggest that a distal p300 binding site contributes to maintenance of promoter-proximal histone acetylation, which promotes gene expression.

Identification of other noncoding elements, such as transcription factor binding sites, that regulate *CUL3* may provide new mechanistic insights into resistance or identify therapeutically tractable targets. To identify candidate transcription factors whose binding sites might be disrupted, we further analyzed via next generation sequencing specific sgRNA target sites after editing and queried these target sites for disruption of known transcription factor motifs using the JASPAR database of transcription factors. At four sgRNA target sites, the canonical transcription factor motifs for Yin Yang 1 (YY1), Zinc Finger Protein 263 (ZNF263), CCCTC-binding factor (CTCF) and activation protein 1 (AP-1) complex were severely disrupted after editing (Fig. 4A) (fig. S5). Based on these observations we hypothesized that mutations within these binding sites abrogate transcription factor recruitment leading to loss of *CUL3* expression and increased vemurafenib resistance. To test these hypotheses, we compared ChIP-ddPCR enrichment of each transcription factor in cells transduced with a sgRNA from our validation set and in control cells (transduced with a non-targeting sgRNA). In the 5´ UTR, two sgRNAs (5´-UTR sg1, sg2) spaced <50 bp apart overlap a YY1 ChIP-seq peak (Fig. 4B). YY1 is a multifunctional transcription factor capable of both gene activation and repression and its overexpression has been observed in various human malignancies (Bushmeyer et al. 1995; Zhang et al. 2011). Analysis of the region using the JASPAR motif and scoring algorithm identifies a canonical YY1 motif with 100% relative score (i.e. the unedited reference sequence perfectly matches the maximum likelihood YY1 motif) (fig. S5A) (Wasserman & Sandelin 2004; Mathelier et al. 2016). After editing with 5´-UTR sg1, the average relative score for the YY1 motif falls to 82% (*n* = 1000 sequencing reads), which is nearly the same as the average score for this motif in random DNA sequences (*n* = 1000 length-matched random sequences) (fig. S5B). Furthermore, we found an increased disruption of the YY1 motif in vemurafenib-treated cells versus vehicle treatment (fig. S6), suggesting that vemurafenib treatment enriches for binding site-damaging mutations. ChIP-ddPCR shows that both sg1 and sg2 decrease YY1 binding, and sg2 (which cuts closer to YY1) more efficiently disrupts YY1 binding than sg1 (67% vs. 26%) (Fig. 4C). In addition, both sg1 and sg2 significantly decrease *CUL3* expression (Fig. 4C). Similarly, 2 sgRNAs in the first intron of *CUL3* (Intron-sg1, sg2) spaced 30 bp apart overlap a ZNF263 ChIP-seq peak (JASPAR relative score: 89%) (Fig. 4D). Both sg1 and sg2 result in a significant decrease in ZNF263 occupancy via ChIP-ddPCR and a decrease in *CUL3* expression (Fig. 4E).

**Figure 4:**
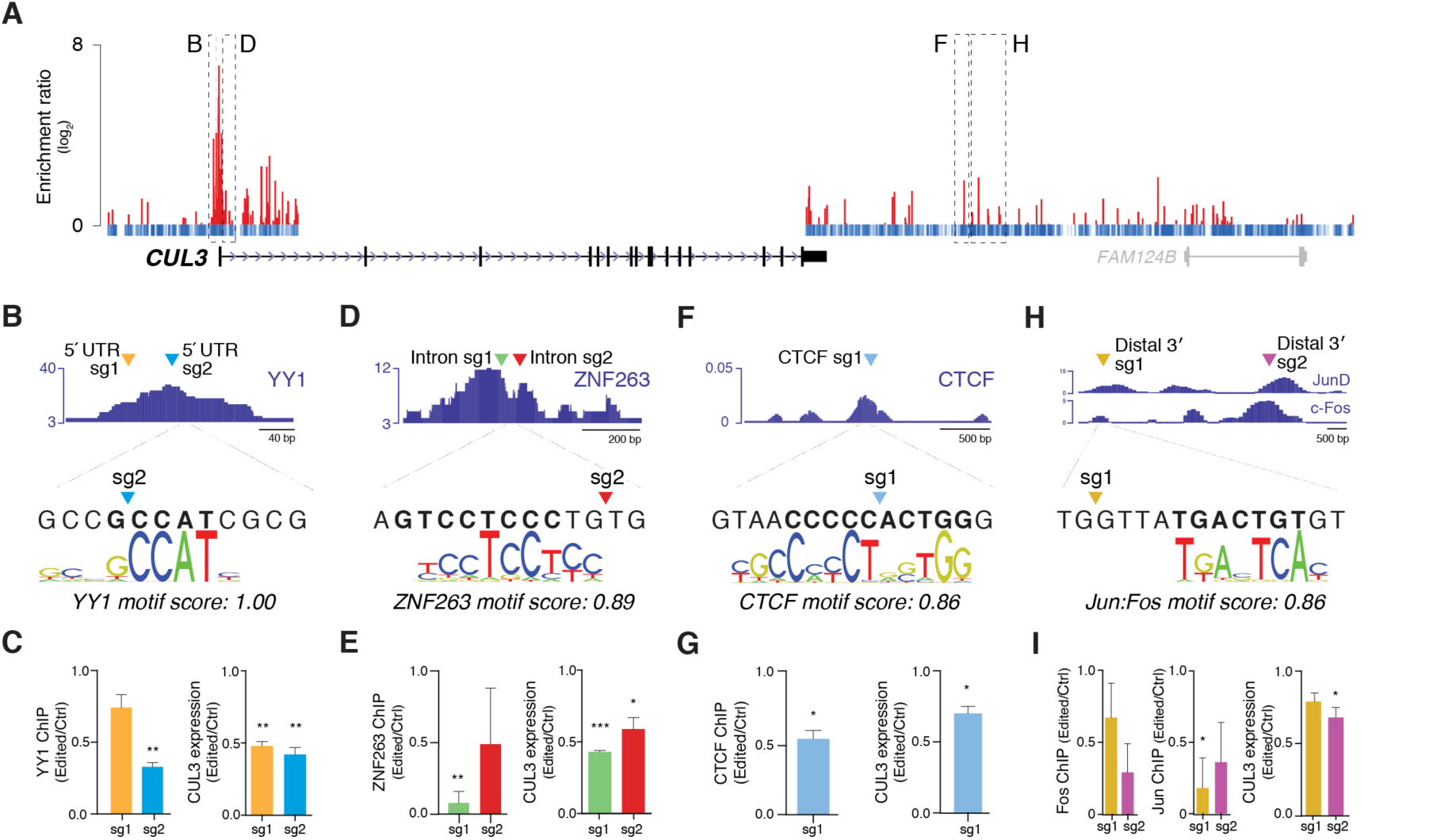
Cas9 mutagenesis disrupts binding of predicted transcription factors and DNA binding proteins at target sites of vemurafenib enriched sgRNAs. A) Location and noncoding screen enrichment of selected sgRNA target sites in the 5’-UTR (*B*,) first intron (*D*) and 3’ distal sites (*F, H*) for transcription factor binding analysis. B)-I) (*top*) Target locations for sgRNAs in relation to bioinformatically-predicted binding sites. Motifs are from the Jaspar vertebrate database and motif scores are Jaspar relative scores (defined as 1 for the maximum-likelihood sequence). ChIP-seq for each region/protein is from K562 cells from ENCODE datasets (SYDH, UChicago OpenChrom/UTAustin). (*bottom*) Change in transcription factor /DNA binding protein occupancy by ChIP around cut site at 7 days post-transduction and change in *CUL3* expression by ddPCR at 7 days post-transduction. Both measurements are normalized to cells transduced with non-targeting sgRNAs.

Although we observe a bias in the presence of regulatory elements 5´ of the transcription start site, we did find several highly enriched sgRNAs downstream of *CUL3*, including two sgRNAs that overlap with AP-1 complex binding sites (distal 3´ sg1, sg2) and another sgRNA that targets a CTCF binding site (CTCF sg1) (Fig. 4F-I). The CTCF sg1 site lies ~30 kb from the 3´ end of *CUL3* and overlaps with non-tissue specific CTCF ChIP-seq peaks of enrichment (Fig. 4F).

CTCF sites are frequently mutated in cancer, and CTCF has been shown to act as an activator, repressor, insulator and mediator of chromatin organization and chromatin loop formation (Katainen et al. 2015; Sanborn et al. 2015). Although we did not find evidence for a strong interaction between this CTCF site and the *CUL3* promoter in our 3C data (~ 0.15 normalized promoter interaction) or in publicly available CTCF chromatin interaction analysis by paired-end tag sequencing (ChIA-PET) (fig. S7), the sgRNA cut site is located in the middle of the predicted CTCF binding motif (JASPAR relative score: 86%). Deep sequencing of the site found mutations in 96% of alleles with a mean indel size (-9.5 bp ± 13.7 bp) that is comparable in size to the canonical CTCF motif. Using ChIP-ddPCR, we found that CTCF occupancy at this site is decreased by 45% after editing and there is a 30% decrease in *CUL3* expression (Fig. 4G). We also explored two putative AP-1 sgRNA target sites that confer drug resistance (Fig. 4H). AP-1 is a heterodimeric basic leucine zipper transcription factor, composed of FOS and JUN subunits, and its over-activation promotes metastasis in carcinomas, breast cancer, and melanoma (Ding et al. 2013). After editing at distal 3´ sg1 and sg2, we found decreased FOS and JUN binding compared with control cells. Editing at either site resulted in an ~25% decrease in *CUL3* expression (Fig. 4I). In keeping with observations in the global screen data, mutation of these 3´ noncoding sites does not have as strong of an effect on gene regulation and function as mutations in the 5´ noncoding region.

Together, our results demonstrate that Cas9-mediated systematic dissection of noncoding loci can identify functional elements involved in gene regulation and altered cancer drug resistance. In combination with other genome-wide assays and datasets, we demonstrate high-throughput identification of regions where changes in chromatin context and transcription factor binding are causally linked to loss of gene expression and a specific, disease-relevant phenotype. This is a generalizable approach, and the extension of pooled CRISPR screens into the noncoding genome will open new inroads into the detection of phenotypically relevant elements and further advance methods for unbiased interrogation of the “Dark Matter” of the genome and its importance in gene regulation.

## Acknowledgements

We would like to thank R. Macrae for critical reading of the manuscript, D. Scott for advice on indel analysis and the entire Zhang laboratory for support and advice. N.E.S. is supported by the NIH through a NHGRI Pathway to Independence Award (K99-HG008171) and a postdoctoral fellowship from the Simons Center for the Social Brain at MIT. J.B.W. is supported by the NIH through a Ruth L. Kirschstein National Research Service Award (F32-DK096822). O.S. is a Klarman Fellow of the Broad-Israel Partnership. F.Z. is a New York Stem Cell Foundation Robertson Investigator and is supported by the NIH through NIMH (5DP1-MH100706 and 1R01-MH110049) and NIDDK (5R01DK097768-03), the New York Stem Cell, Simons, Paul G. Allen Family, and Vallee Foundations; and David R. Cheng, Tom Harriman, and Robert Metcalfe. Reagents are available to the academic community through Addgene and associated protocols, support forums, and computational tools are available via the Zhang lab website (www.genome-engineering.org).

